# Programmable marine bacteria catalyze the valorization of lignin monomers

**DOI:** 10.1101/2024.03.12.584725

**Authors:** Ying Wei, Shu-Guang Wang, Peng-Fei Xia

**Author notes:** Correspondence: Peng-Fei Xia, School of Environmental Science and Engineering, Shandong University, Qingdao 266237, China.

## Abstract

Efficiently converting lignin, the second most abundant biopolymer on Earth, into valuable chemicals is pivotal for a circular economy and net-zero future. However, lignin is recalcitrant to bio-upcycling, demanding innovative solutions. We report here the biological valorization of lignin-derived aromatic carbon to value-added chemicals without requesting extra organic carbon and freshwater via reprogramming the marine *Roseobacter* clade bacterium *Roseovarius nubinhibens*. We discovered the unusual catalytic advantages of this strain for the oxidation of lignin monomers and implemented a CRISPR interference (CRISPRi) system with the *lacI*-P_trc_ inducible module, nuclease-deactivated Cas9, and programmable gRNAs. This enabled precise and efficient repression of target genes. By deploying the customized CRISPRi, we reprogrammed the carbon flux from a lignin monomer, 4-hydroxybenzoate, to achieve maximum production of protocatechuate, a pharmaceutical compound, while maintaining essential carbon for cell growth and biocatalysis. As a result, we achieved a 4.89-fold increase in protocatechuate yield with a dual-targeting CRISPRi system. Our study introduces a new-to-the-field lineage of marine bacteria and underscores the potential of blue biotechnology leveraging resources from the ocean for simultaneous carbon and water conservation.

## Introduction

Reverting carbon losing to the environment back to resources for carbon-conservative bioproduction is an important route towards a net-zero future. Lignin, a prevalent aromatic polymer, represents a substantial fraction, typically ranging from 15% to 40%, of the dry mass of terrestrial plants on Earth (1, 2). However, lignin is largely underutilized due to its inherent recalcitrance to biodegradation and heterogeneity in composition (3–6). Consequently, lignin-containing biomass (e.g., plant cell walls and agriculture wastes) often demands disposal via, for instance, incineration, causing CO_2_ emission and environmental pollution. Innovations for the valorization of lignin are imperative and urgent. Given the revolutionary advances in engineering biology, bioconversion of lignin-derived aromatic compounds to value-added chemicals, e.g., *cis*,*cis*-muconic acids, pyridine-dicarboxylic acids and β-ketoadipic acids, has been achieved in engineered *Pseudomonas putida* (7–9), *Corynebacterium glutamicum* (10) and *Rhodococcus jostii* (11), illustrating a promising avenue for the sustainable valorization of lignin-derived aromatics.

One often overlooked sector, however, is the source of water to sustain these enabling biotechnologies for, but not limited to, lignin valorization. A large amount of freshwater is commonly necessary to cultivate engineered microbes (12, 13), competing with human needs for freshwater. This will be a controversial issue, especially with the fact of global water scarcity (14–16). To tackle this conflict, we propose blue solutions by leveraging marine bacteria and resources from the ocean (17). Marine bacteria usually survive or even thrive in environments with limited nutrients, extreme temperatures and high salinity, which endow them with versatile metabolic capabilities and tolerances to various stressors (18–20). Besides, marine bacteria inherently grow in seawater avoiding direct competition for freshwater resources with humans. Therefore, marine bacteria can be engineered as promising microbial chassis for lignin bio-valorization with seawater, resolving the challenges in carbon and water conservation simultaneously.

*Roseovarius nubinhibens* is a member of the marine *Roseobacter* clade with diverse metabolic capacities, notably including the degradation of aromatics through β-ketoadipate pathway (21). The natural properties enable *R. nubinhibens* a potential whole-cell catalyst for the conversion of lignin-derived aromatic compounds into value-added chemicals. Though bacteria in this clade have been intensively interrogated for the ecological and evolutionary dynamics (22–24), the potential of these bacteria for biocatalysis has rarely been discovered. Notably, genetic systems are available for strain engineering, including the delivery of foreign DNAs, replicable shuttle vectors, and “knock-in” gene disruption (25–27). In a previous work, we established a CRISPR-Cas-based genome editing tool at a single-nucleotide resolution for *R. nubinhibens*, and efficient and precise gene inactivation can be realized, paving the way for the design and construction of strains capable of lignin upcycling (28).

Here, we report the biological valorization of lignin monomers catalyzed by the marine bacterium *R. nubinhibens* with a multiplex CRISPR interference (CRISPRi) system. We reveal the unusual advantages of this strain in catalyzing the oxidation of lignin monomers, underscoring its potential as a versatile biocatalyst. We modularly designed the CRISPRi system with nuclease-deactivated Cas9 (dCas9) from *Streptococcus pyogenes* and programmable gRNAs driven by an inducible system. Then, we selected 4-hydroxybenzoate (4HB) as a proof-of-principle lignin monomer and protocatechuate (PCA), a pharmaceutical compound, as the product to evaluate the bioproduction by *R. nubinhibens* with CRISPRi. We present here a new-to-the-field lineage of marine bacteria as whole-cell biocatalyst for lignin valorization, and a paradigm of leveraging marine bacteria with cutting-edge synthetic biology tools for sustainable biorefinery. Beyond carbon, we are appealing for more attention to the water requirement of bio-innovations to ensure a sustainable future.

## Results

### PCA accumulation during cell growth on 4-hydroxybenzoate

**Fig. 1.**
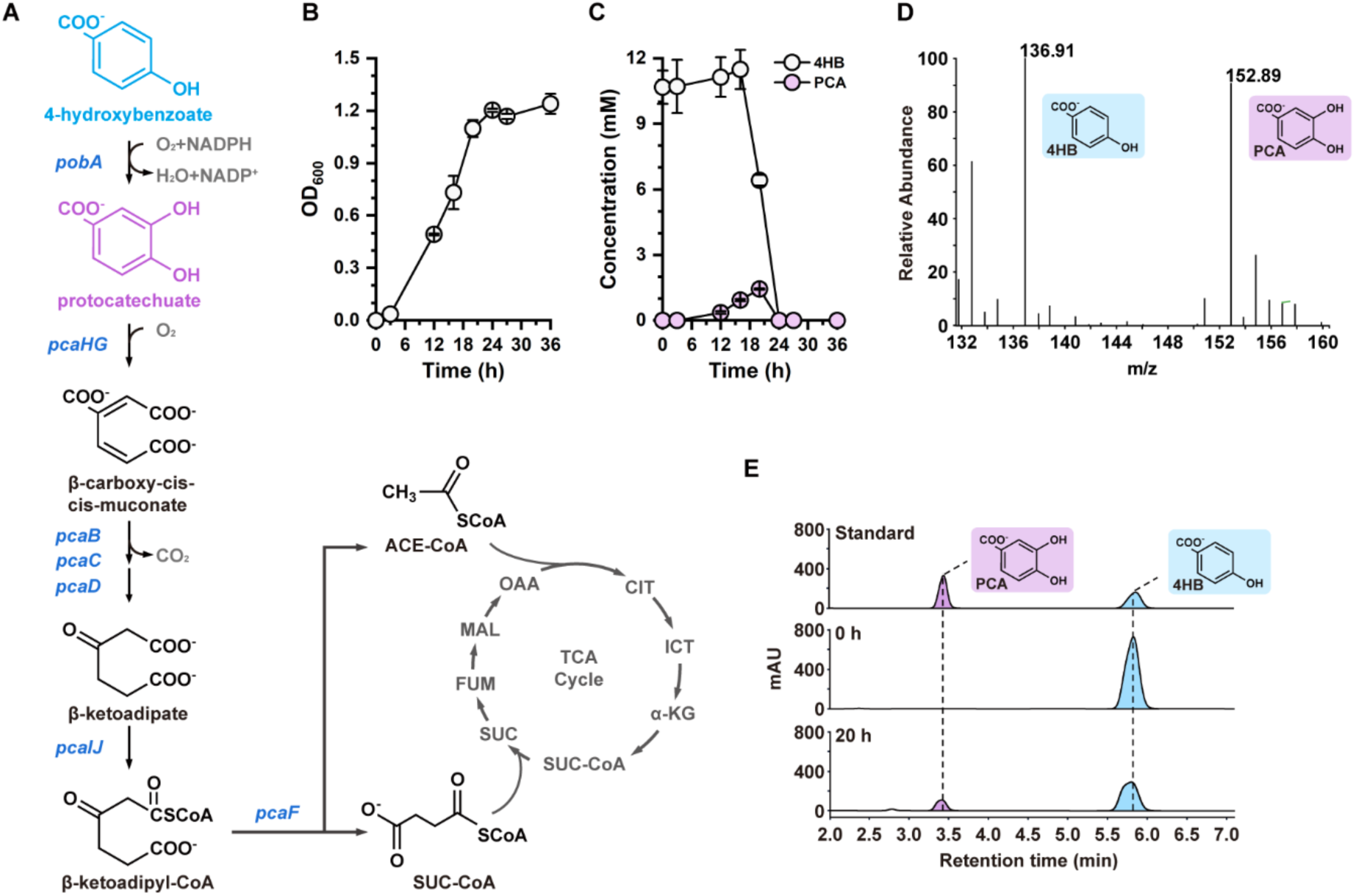
PCA accumulation in *R. nubinhibens*. **(A)** The protocatechuate branch of β-ketoadipate metabolic pathway in *R. nubinhibens.* CIT, citrate; ICT, isocitrate; α-KG, alpha-ketoglutarate; SUC-CoA, succinyl-CoA; SUC, succinate; FUM, fumarate; MAL, malate; OAA, oxaloacetate; ACE-CoA, acetyl-CoA. **(B)** Growth profile. **(C)** Utilization of 4HB and accumulation of PCA in the wild-type *R. nubinhibens* with 4HB as the sole carbon source. Experiments were carried out in triplicate and the error bars represented the standard deviations of the means of three biological replicates. **(D)** MS analysis of samples. **(E)** HPLC profile of standard and samples.

The protocatechuate branch of the β-ketoadipate pathway is the main pathway for the metabolism of lignin-derived aromatic molecules (29, 30). Going through this pathway, the aromatic carbon goes to the tricarboxylic acid (TCA) cycle with two key metabolites, succinyl-CoA and acetyl-CoA, as the nodes, providing carbon and energy for the anabolisms and catabolism in the cell (**Fig. 1A**). To evaluate the potentials of *R. nubinhibens* for lignin degradation, we selected 4HB, a representative and readily available lignin-derived aromatic compound monomer, as the substrate and grew the strain in a marine minimal medium. Notably, *R. nubinhibens* demonstrated robust growth with 4HB as the sole carbon source without extra organic substrates, such as glucose, to power the metabolism (**Fig. 1B and 1C**). Interestingly, during the cultivation of the wild-type *R. nubinhibens* in the basal medium with 4HB, a gradual accumulation of a violet compound was observed with the growth of cells in the first 20 h **(Fig. S1)**. Subsequent characterization via MS and HPLC confirmed the identity of this compound as PCA (**Fig. 1D and 1E and Fig. S2)**, a pharmaceutical compound and central metabolite in the β-ketoadipate pathway. PCA can also be a platform compound for the production of diverse value-added chemicals, such as *cis*,*cis*-muconic acid, adipic acid and levulinic acid (31–33). However, conventional PCA production relies on extraction from plants with low yield and high cost (34). This observation implied an unusually high catalytic capability of the native 4-hydroxybenzoate 3-monooxygenase, PobA, for 4HB hydroxylation to PCA, which is usually a limiting step for aromatic utilization (35–37). Therefore, a rational allocation of carbon flux from 4HB would be possible to maximize the biosynthesis of PCA and to sustain cell growth simultaneously. This can be realized using a CRISPRi system as a carbon-flux-limiting stopcock to restrain the flux towards the TCA cycle by downregulating relevant genes downstream of *pobA* (**Fig. 1A**).

### Characterization of a *lacI*-P_trc_ inducible system in *R. nubinhibens*

One of the key advantages of CRISPRi is that the repression of target genes is tunable, which usually relies on an inducible system based on transcriptional regulators. Currently, no inducible system has been developed for the *Roseobacter* clade bacteria. As such, we designed and constructed a *lacI*-P_trc_ inducible system consisting of the *lacI* gene coding for the repressor LacI and the corresponding promoter P_trc_ driving the expression of red fluorescent protein, mCherry, as a reporter. Without inducers, LacI would bind to the operator region of the P_trc_ promoter blocking the expression of mCherry, while, when IPTG is supplemented, the repressor would ligand to IPTG instead and release the promoter for the transcription of mCherry (**Fig. 2A**).

To assess the functionality of the *lacI*-P_trc_ inducible system, we monitored the fluorescence of *R. nubinhibens* with pmCherry **(Table S1)** and the control plasmid (pBBR1MCS-5, **Table S1**) via fluorimetry and flow cytometry. Without IPTG, both strains showed a similarly low level of background fluorescence. We tested eight different concentrations of IPTG from 0.01 mM to 2 mM. The strain displayed a gradually increased signal of fluorescence along with the rising concentration of IPTG from 0.01 mM to 0.5 mM, and the signals indicated a saturation of IPTG induction when the concentration reached 0.5 mM (**Fig. 2B**). The proportion of fluorescent cells reached saturation at 29.9% with 0.5 mM of IPTG (**Fig. 2C and 2D**). This is the first demonstration of an inducible system in *R. nubinhibens*, making dynamic regulation of designed genetic modules possible.

**Fig. 2.**
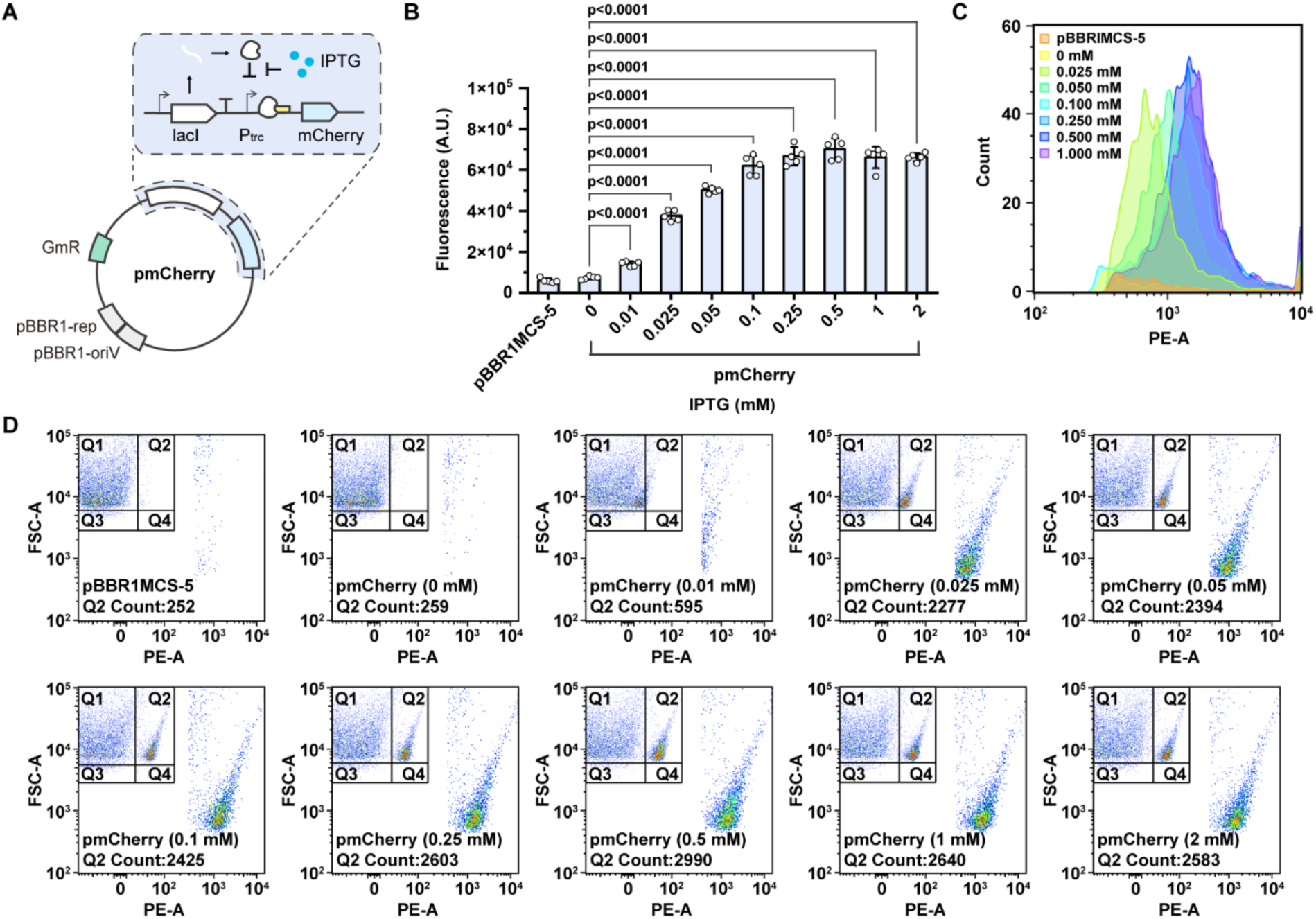
The *lacI*-Ptrc inducible system for *R. nubinhibens*. **(A)** Schematic representation of the pmCherry. The *lacI* gene is driven by a constitutive promoter and the mCherry fluorescence gene is driven by the Ptrc inducible promoter. **(B)** Fluorescence of strains with pBBR1MCS-5, as a control, and pmCherry under different IPTG concentrations. The value of fluorescence intensity (A. U., Arbitrary Unit) was normalized by optical density at 600 nm. Experiments were carried out in quintuplicate, and the error bars represented the standard deviations of the means of five biological replicates. The differences were statistically evaluated by t-test. **(C)** Count of mCherry fluorescent cells. **(D)** Distribution of mCherry fluorescent cells in the population. For each sample, 10,000 events were analyzed.

### Modular design of a CRISPR interference system for *R. nubinhibens*

To implement a CRISPRi system in *R. nubinhibens*, we modularly assembled the inducible module based on the *lacI*-P_trc_ system and an interference module containing *dcas9* and the gRNA cassette, carried by the pBBR1MCS-5 vector plasmid (**Fig. 3A**). In the design, dCas9 driven by the *lacI*-P_trc_ system reserves the ability to bind the target DNA (38–40), and navigated by a programmable gRNA, it binds to the target and sterically blocks RNA polymerase (RNAP), thereby interfering the transcription process and eventually repressing the expression of the target gene (**Fig. 3B**) (41–43). The gRNA cassette can be programmed for any genes of interest by substitution of the 20-nt spacer via a one-step PCR protocol (**Fig. 3C**).

**Fig. 3.**
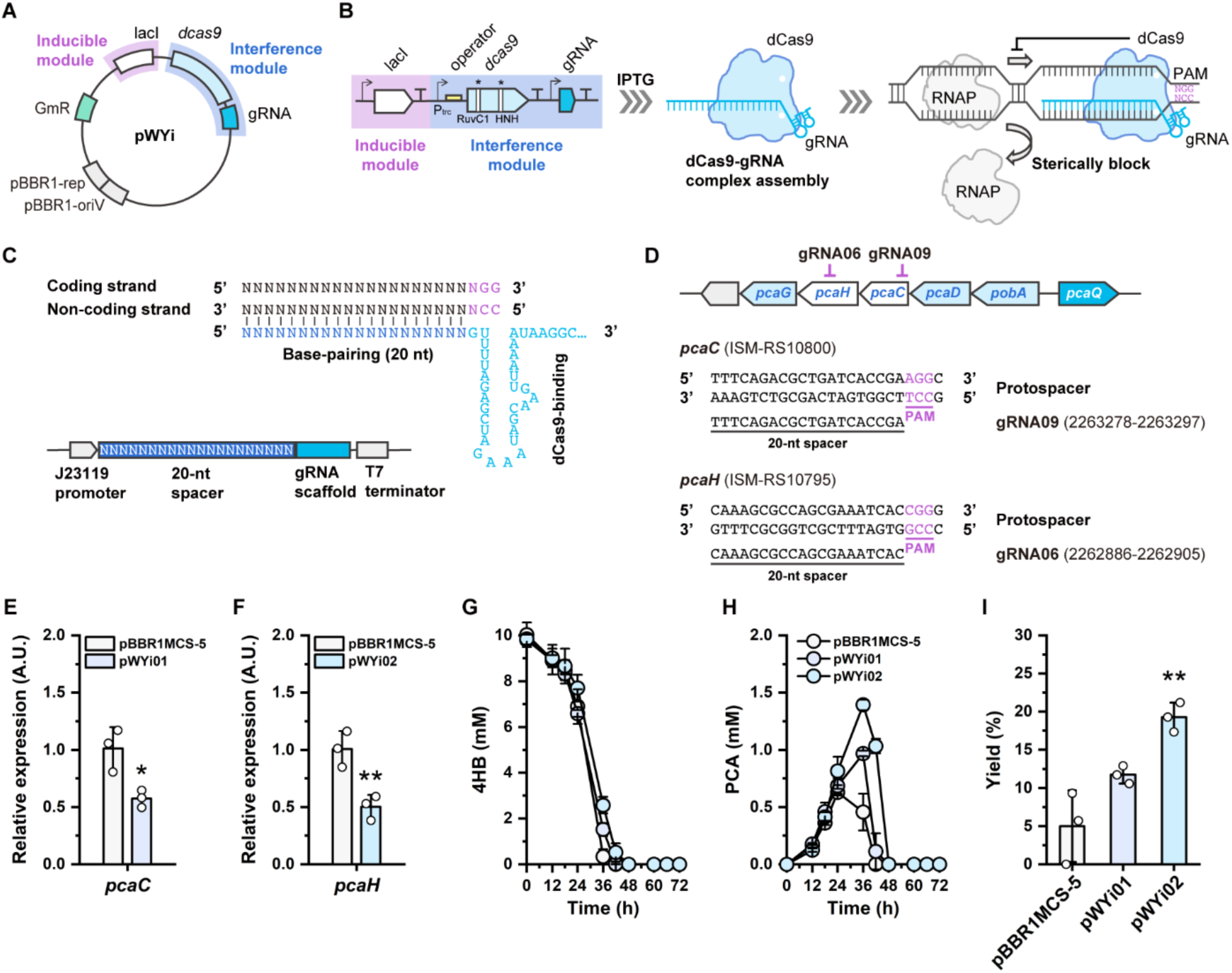
CRISPRi system for *R. nubinhibens*. **(A)** Design of the working plasmid pWYi. The pWYi plasmid was comprised of the inducible module (*lacI*-Ptrc inducible system) and the CRISPR interference module (*dcas9* and the gRNA cassette). **(B)** Schematic representation of the working principle of CRISPRi. The gRNA and dCas9 (D10A and H840A) complex bind to the target DNA and sterically block the RNAP. The site of protospacer adjacent motif (PAM) was highlighted in violet. **(C)** Structure of the gRNA cassette targeting the coding strand. The sequences of the 20-nt base-pairing region of the gRNA are identical to the coding strand. The base-pairing region was indicated in navy blue and the dCas9 binding region was indicated in sky blue. **(D)** Positions of the gRNAs targeting the *pca* gene cluster. **(E)** Relative expression of target genes in the strain with pWYi01 and **(F)** pWYi02. **(G)** Utilization of 4HB. **(H)** Titer of PCA. **(I)** Molar yield (%) of PCA. Samples were taken at the exponential phase with 0.25 mM and 0.5 mM IPTG and the strain with pBBR1MCS-5 was included as the control. Experiments were carried in triplicate and the error bars represented the standard deviations of the means of three biological replicates. The differences were statistically evaluated by t-test (*, p<0.05; **, p<0.01).

As a proof of concept, we attempted to target two genes, *pcaC* and *pcaH,* in the genome, respectively. Both genes are involved in the β-ketoadipate pathway, which has been identified as the primary route to degrade aromatic compounds in *R. nubinhibens* (21). Specifically, the *pcaC* gene encodes the γ-carboxymuconolactone decarboxylase and *pcaH* encodes the protocatechuate 3,4-dioxygenase β-subunit catalyzing the ring cleavage together with PcaG (**Fig. 1A**). We constructed two working plasmids, pWYi01 (with gRNA09, **Table S1**) and pWYi02 (with gRNA06, **Table S1**) targeting *pcaC* and *pcaH*, respectively (**Fig. 3D**). IPTG (0.25 mM and 0.5 mM) were added to induce the expression of dCas9, and the expression of target genes was determined by RT-qPCR. As expected, the expression of *pcaC* and *pcaH* in strains with pWYi01 or pWYi02 significantly decreased by 43.13% and 50.26%, respectively, compared to strains with the control plasmid under 0.5 mM IPTG induction (**Fig. 3E and 3F**). The repression degree with 0.5 mM IPTG was, though marginal, higher than that with 0.25 mM IPTG, implying a higher expression level of dCas9 which is in agreement with our characterization above **(Fig. S3)**. These results demonstrated that the CRISPRi system was successfully established.

The *pcaH* and *pcaC* genes, that we targeted with CRISPRi thereof, are located downstream of *pobA* in the *pca* gene cluster. The repression of these downstream genes is expected to promote the accumulation of upstream metabolites (**Fig. 3D and 1A**). So, we tested the strains with the CRISPRi system targeting *pcaH* and *pcaC*, respectively, to evaluate the conversion of 4HB to PCA. Both strains were cultivated in shake flasks with 4HB as the sole substrate and induced with 0.5 mM IPTG. Compared with the control, both strains with CRISPRi showed slower growth rates and 4HB consumption (**Fig. 3G and Fig. S4)**. The strain with the control plasmid pBBR1MCS-5 produced PCA at a titer of 0.46 ± 0.43 mM and a molar yield of 5.00 ± 4.69% at 36 h (**Fig. 3H and 3I and Table S2)**. As expected, the strains with CRISPRi displayed higher performance with improved titer and yield of PCA. The strain expressing pWYi02 (targeting *pcaH*) exhibited a 3.85-fold increase in PCA yield, while that of the strain expressing pWYi01 (targeting *pcaC*) increased by 2.35-fold than the control strain (**Fig. 3I**). Specifically, the strain with CRISPRi repressing *pcaH* and *pcaC* produced 1.39 ± 0.06 mM and 0.97 ± 0.03 mM of PCA with a molar yield of 19.27 ± 1.93% and 11.74 ± 1.16% at 36 h, respectively (**Fig. 3H and 3I and Table S2)**. Under this condition, strains with 0.25 mM and 0.5 mM IPTG showed similar performance in the growth rate **(Fig. S5)**, 4HB consumption and accumulation of PCA **(Fig. S6 and Fig. S7)**, indicating that the difference between these two inducible strengths is marginal. These results demonstrated that the CRISPRi system efficiently allocated the carbon flux and could be modified as a powerful tool for the dynamical regulation of bioproduction in *R. nubinhibens*. However, the CRISPRi system with only one target was not sufficient, and further improvement is necessary.

### Multiplex CRISPRi enables enhanced biosynthesis of PCA

To further enhance the accumulation of PCA, we developed a multiplex CRISPRi system with tandem gRNA cassettes targeting different genes in the operon simultaneously. We selected *pcaG*, encoding the protocatechuate 3,4-dioxygenase α-subunit, which directly catalyzes the PCA ring-opening process together with PcaH, as another target. The tandem gRNA modules were generated by assembling two gRNA cassettes (**Fig. 4A**). The resulting working plasmids pWYi-M03 (with gRNA09 and gRNA06, **Table S1**) and pWYi-M04 (with gRNA06 and gRNA07, **Table S1**) target *pcaC* and *pcaH* or *pcaH* and *pcaG*, respectively (**Fig. 4B**). We first evaluated the expression of the target genes at the transcriptional level. As intended, all strains with CRISPRi exhibited downregulated expression of the target genes. The expression of strains harboring the multiplex CRISPRi system was lower than that of strains with a single target (**Fig. 4C and 4D**). The strain with pWYi-M03 descended the expression of *pcaC* and *pcaH* by 22.38% and 24.47%, respectively, compared to the strain with one gRNA (**Fig. 4C**). Multiplex CRISPRi system regulating the *pcaHG* genes displayed the strongest interference and declined the expression of *pcaH* and *pcaG* by 57.80% and 69.69% than the control, respectively (**Fig. 4D**). These data indicated the stronger interference of genes at the RNA level with multiple locus targeting than single sites.

**Fig. 4.**
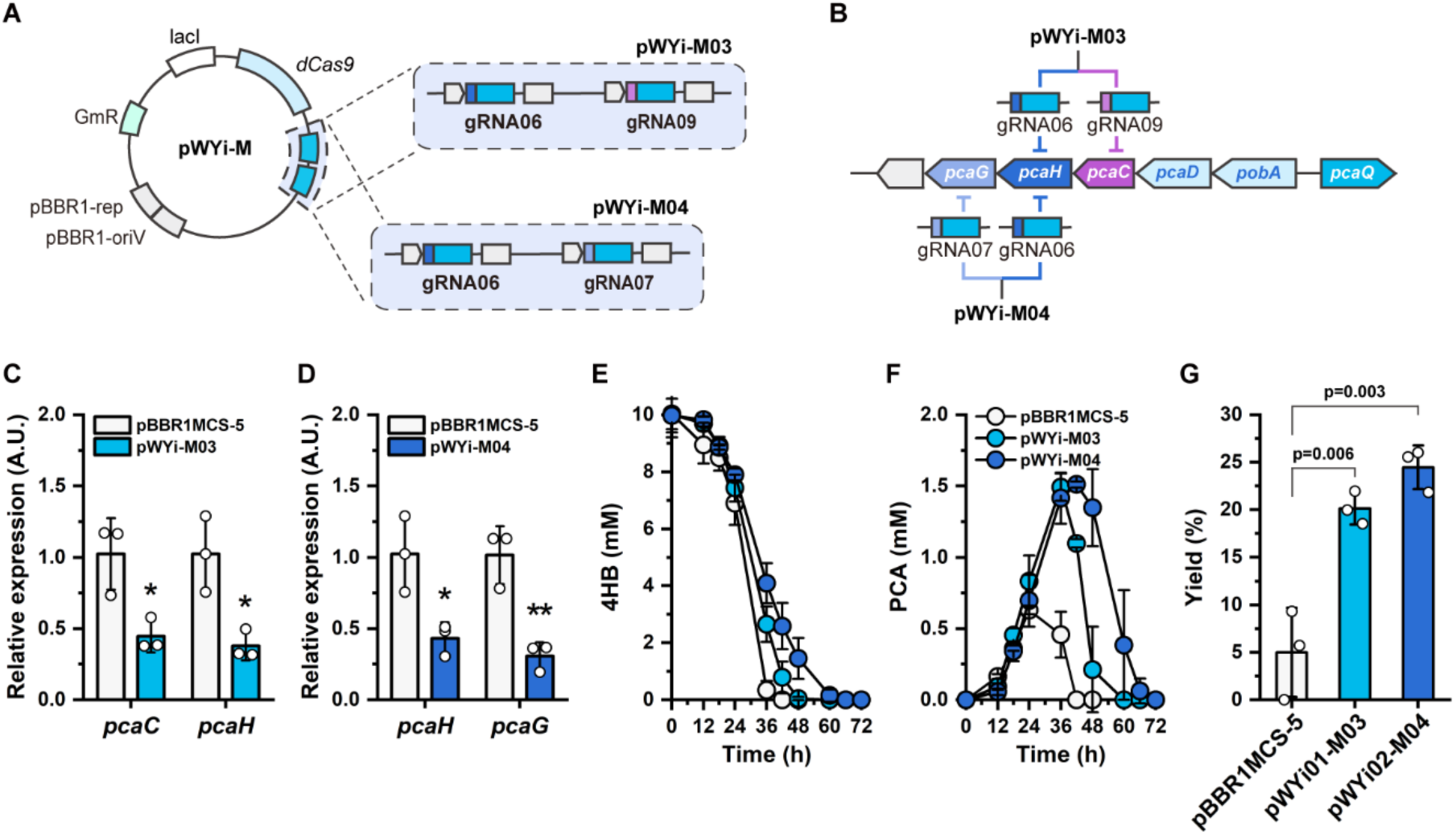
Multiplex CRISPRi for *R. nubinhibens*. **(A)** Design of pWYi-M. The multiple gRNAs module consisted of two tandem gRNA cassettes. **(B)** Positions of gRNAs targeting in the *pca* gene cluster. **(C)** Relative expression of target genes in the strain with pWYi-M03, and **(D)** pWYi-M04. **(E)** Utilization of 4HB. **(F)** Titer of PCA. **(G)** Molar yield (%) of PCA. Samples were taken at the exponential phase with 0.5 mM IPTG induction and the strain with pBBR1MCS-5 was included as the control. Experiments were carried out in triplicate and the error bars represented the standard deviations of the means of three biological replicates. The differences were statistically evaluated by t-test.

With the multiplex CRISPRi system, we found that the growth of strains with all designed CRISPRi systems was repressed, while the multiplex CRISPRi targeting *pcaHG* resulted in the lowest growth rate **(Fig. S8)**. Consistent with the growth rate, multiplex CRISPRi showed stronger interference in the utilization of 4HB than single-targeting CRISPRi (**Fig. 4E and Table S3)**. PCA production was raised in strain with pWYi-M03 (targeting *pcaC* and *pcaH*) at a titer of 1.49 ± 0.12 mM at 36 h, an increase of 1.54-fold and 1.07-fold over the strain with pWYi01 and pWYi02, respectively (**Fig. 4F and Table S3)**. The maximum titer of PCA appeared in the strain harboring pWYi-M04 (targeting *pcaH* and *pcaG*) with 1.51 ± 0.03 mM at 42 h and raised 13.45-fold and 1.46-fold than the strain individually targeting *pcaC* and *pcaH*, respectively (**Fig. 4F and Table S3)**. We calculated the molar yields of each strain at 36 hours, and the strain with pWYi-M04 exhibited the highest yield at 24.47 ± 2.30%, a 2.08-fold and 1.27-fold increase over the strain with pWYi01 and pWYi02, respectively (**Fig. 4G and Table S3)**. Yet, little improvement of the yield was observed in the strain with pWYi-M03 repressing *pcaC* and *pcaH* when compared to the strain only repressing *pcaH* alone **(Table S2 and Table S3)**. This can be explained by analyzing the β-ketoadipate pathway. PCA was catalyzed into β-carboxy-*cis*-*cis*-muconate by PcaG and PcaH. As a result, interference of these enzymes, compared to PcaC, would directly impede the aromatic ring-cleavage, therefore promoting the accumulation of PCA. While the strain with the control plasmid only produced PCA with the maximum yield at 5.00 ± 4.69% at 36 h, the strain with pWYi-M04 showed an increase of 4.89-fold than the control (**Fig. 4G and Table S3)**. To be noticed, we also observed that PCA was retained in the culture for 72 h and prolonged nearly double the accumulation time, providing biosynthesis with an enlarged operation window to control the reaction dynamics or terminate the process at the maximal titer of bioproducts.

## Discussion

In this study, we report the biological valorization of lignin-derived aromatic carbon into value-added chemicals without requesting extra organic carbon and freshwater via reprogramming the metabolism of the marine bacterium *R. nubinhibens*. Our findings highlight the unique advantages offered by *Roseobacter* clade bacteria, particularly in the context of lignin valorization. An obvious benefit is the high catalytical activity of PobA in *R. nubinhibens*, which is no longer a bottleneck but leads to the accumulation of PCA in the β-ketoadipate pathway (**Fig. 1C**). To uncover the potential of *R. nubinhibens*, we established a *lacI*-P_trc_ inducible system and developed a CRISPRi system for the repression of genes of interest in *R. nubinhibens*. We successfully targeted the key genes (*pcaC*, *pcaH* and *pcaG*) in the β-ketoadipate pathway individually or in tandem, generating a carbon-flux-limiting stopcock for rational carbon re-allocation. By doing so, we maximized the carbon accumulation before the ring-cleavage step as PCA and allowed minimal flux going to the TCA cycle to sustain cell growth and provide energy for biocatalysis. When targeting *pcaH* and *pcaG* simultaneously, the conversion of 4HB to PCA exhibited a 4.89-fold increase in molar yield (**Fig. 4G**), showing a great potential of the blue bio-valorization of lignin monomers via the dynamic regulation of marine bacteria with multiplex CRISPRi. Nevertheless, we acknowledge that the accumulated PCA was eventually metabolized for cell growth if it was not harvested in a certain time frame. Even though our CRISPRi system expanded the operational window to around 36 - 48 hours without sacrificing maximum yield and titer. A combination of gene regulation and editing would be beneficial for further advances, and *R. nubinhibens* may be further equipped with designed new features (e.g., unnatural substrates or products) for enhanced upcycling of lignin monomers.

Upcycling carbon losing to the environment is vital for a net-zero future. Lignin-rich biomass commonly needs conventional disposal via incineration, leading to considerable carbon loss and air pollution, for instance, the emission of greenhouse gases CO_2_, NO_x_ and inhalable particles (e.g., PM10 and PM2.5) (44–46). In contrast, biological valorization has been considered a carbon-efficient, economically viable and environmentally friendly alternative to upcycle lignin to high-performance bioproducts. Two long-standing but often overlooked issues in, but not limited to, lignin bio-valorization remain. One is the requirement of extra organic carbon sources, e.g., glucose, during fermentation (10, 35, 47), aggravating the “food-versus-fuel” debate. The other is the demand for water to cultivate engineered microbes, potentially rousing the competition of freshwater between humans and biosynthesis, especially under global water scarcity (48–50). Inspiringly, the engineered *R. nubinhibens*, to some extent, provides a solution to tackle these issues at once. First, this strain can metabolize 4HB as a sole carbon source without demanding any extra carbon, and the reprogrammed carbon flux can not only support cell growth but also drive the biocatalysis of 4HB to PCA. Second, *R. nubinhibens*, as a marine bacterium, grows in the ocean with natural tolerance to high salinity, making it possible and preferable to use seawater for cultivation and fermentation. We believe that blue bio-valorization by engineered marine microbes will be a new wave for building a circular economy and contributing to multiple Sustainable Development Goals (SDGs) related to the climate, carbon and water, such as SDG6 (Clean water and sanitation), SDG12 (Responsible consumption and production), and SDG13 (Climate action). Beyond these, blue bio-valorization depending on seawater rather than the unequally distributed freshwater may also contribute to reducing inequality (SDG10) in the wave of biotechnological revolution.

## Methods

### Strains and media

*E. coli* DH5α (Takara Bio) was used for molecular cloning and cultivated in Luria-Bertani (LB) liquid media containing 10 g NaCl, 10 g tryptone, and 5 g yeast extract per liter or on LB solid agar (1.5%) plates at 37 °C. When screening *E. coli* transformants and maintaining plasmids, appropriate antibiotics were added (100 µg mL^-1^ ampicillin and 20 µg mL^-1^ gentamicin). *R. nubinhibens* ISM (DSM 15170) was grown at 30 °C and 180 rpm in Difco^TM^ Marine Broth 2216 (MB2216) liquid media or on solid plates for general cultivation, and in the marine basal medium containing 200 mM NaCl, 50 mM MgSO_4_, 10 mM CaCl_2_, 10 mM KCl, 10 mM NH_4_Cl, 1 mM K_2_HPO_4_, 0.1 mM FeEDTA, 0.05% yeast extract and 0.1% vitamin with 10 mM 4HB as the carbon source for biosynthesis. 20 µg mL^-1^ of gentamicin was used for isolating clones of *R. nubinhibens*.

### Plasmid construction and electroporation

All plasmids used in this study were summarized in **Table S1**, and all primers used in this study were summarized in **Table S4**. All the gRNA cassettes were summarized in **Table S5**. The synthesized sequence of mCherry was listed in **Table S6**. The mCherry fragment carried by pQLL plasmid with Amp^R^ was synthesized by Beijing Liuhe BGI and then amplified using PrimeSTAR^®^ Max DNA Polymerase (Takara Bio). The plasmid pmCherry was generated by fusing the *lacI*-P_trc_ inducible system and mCherry into the vector pBBR1MCS-5 using In-Fusion^®^ Snap Assembly Master Mix (Takara Bio). All plasmids were extracted using QIAprep^®^ Spin Miniprep Kit (Qiagen). The gRNA09 cassette targeting *pcaC* was built via inverse PCR using back-to-back primers containing the 20-bp spacer. Based on the plasmid pWY06 (28), we constructed pWYi02 by eliminating the deaminase, and then pWYi01 was constructed by substituting the gRNA cassette. To generate multiplex CRISPRi working plasmids, the gRNA09 and gRNA07 cassettes were assembled in the pWYi02, forming pWYi-M03 and pWYi-M03, respectively.

For the transformation of working plasmids, the electrocompetent cells were prepared by washing twice using the buffered sucrose solution and stored at -80 °C before use (28). The working plasmids were added to 80 µL competent cells in a pre-cooled 0.1-cm MicroPulser Electroporation Cuvette (Bio-Rad) and the mixture was treated in MicroPulser Electroporator (Bio-Rad) at the pulse intensity of 1.8 kV and resulting pulse length of 1.3-1.5 ms. Cells after electroporation were transferred to fresh media for recovery and then plated on a solid medium supplemented with 20 µg mL^-1^ gentamicin.

### Fluorimetry and flow cytometry

The strains transformed with pBBRIMCS-5 and pmCherry were cultivated overnight in MB2216 medium with 20 µg mL^-1^ gentamicin. Next, aliquots of 2-mL suspensions were induced for 4 h with IPTG at different concentrations from 0 mM to 2 mM. After induction, the cells were harvested and washed twice using phosphate-buffered saline (PBS). For the fluorimetry assay, the resuspended cells were diluted 10^2^ times, and 200 µL of each bacterial suspension was transferred to 96-well microplates (Corning Costar). The OD_600_ and fluorescence intensity at an excitation wavelength of 590 nm and an emission wavelength of 645 nm were determined using Spark^®^ Multimode Microplate Reader (TECAN). The fluorescence intensity (Arbitrary Unit) was normalized for fair comparation. For flow cytometry, the resuspended cells were diluted by 10^3^ times and then stored at 4 °C until analysis. The mCherry fluorescence distribution at the population level was analyzed using BD FACS Aria Fusion (BD Biosciences) equipped with a 561-nm laser. 10,000 single-cell events were collected for each sample. Data were processed using FlowJo (TreeStar Inc.).

### Cultivation experiment

For biosynthesis, the strains from glycerol stocks were cultivated in MB2216 media overnight with appropriate antibiotics. Next, the culture was inoculated into 250 mL shake flasks containing 50 mL fresh marine basal medium with 4HB (10 mM) as the carbon source by 1:50 dilution and cultivated at 30 °C and 180 rpm. Gentamicin (20 µg mL^-1^) was added to maintain the working plasmids and IPTG (0.25 mM or 0.5 mM) was supplemented for induction. Samples were taken at different intervals and stored at -20 °C until analysis. OD_600_ was measured to determine the cell growth profiles of each sample. The concentration of 4HB and PCA was quantified using HPLC as described below. At least two parallel biological experiments for each strain were performed and the mean value with standard deviation was calculated and reported.

### Quantification of mRNA expression levels

After induction for 24 h, cells with the CRISPRi systems were harvested by centrifugation at 10,000 rpm and frozen with liquid nitrogen, and then stored at -80 °C until extraction. Total RNA was extracted using RNAprep Pure Cell/Bacteria Kit (TIANGEN) and reverse-transcribed into cDNA using PrimeScript™ RT reagent Kit with gDNA Eraser (Perfect Real Time) (Takara Bio). The resulting cDNA was utilized for analysis using TB Green^®^ Premix Ex Taq™ II (Tli RNaseH Plus) (Takara Bio). Quantitative PCR was performed using Applied Biosystems QuantStudio™ 5 (Thermo Fisher Scientific). To normalize the gene expression between different samples, the gene encoding for the 16S ribosomal RNA was employed as the housekeeping gene. The primers used for RT-qPCR were listed in **Table S4**.

### Identification and quantification of 4HB and PCA

Samples were centrifuged at 10,000 rpm for 1 min and the supernatants were sterilely filtered by 0.22-µm filter and diluted by 10 times. For identification of products, samples were demineralized by solid phase extraction using HyperSep™ C18 (Thermo Scientific) and analyzed using HPLC-MS (LCQ Fleet, Thermo Fisher Scientific). For quantification of 4HB and PCA, samples were detected by Agilent 1260 HPLC equipped with a UV detector (Agilent Technologies) at a wavelength of 210 nm with an injection volume of 10 µL. Chromatographic separation was realized by an EC-C18 column 4 µm, 4.6 × 100 µm (Agilent Technologies) at 25 °C with the mobile phase comprised of 90% formic acid (0.1%) and 10% acetonitrile at a constant flow rate of 0.8 mL per min. Molar yield (%) of PCA was calculated by dividing the titers of PCA by 4HB.

## Conflict of Interests

The authors declare no conflict of interest.

## Acknowledgments

The authors thank Dr. Haiyan Yu for the assistance with flow cytometry analysis and Dr. Fanping Zhu for the help with HPLC-MS analysis. This work was supported by the National Natural Science Foundation of China (22278246, U20A20146 and 22378233), the Department of Science and Technology of Shandong Province (2022HWYQ-017), the Natural Science Foundation of Shandong Province (ZR2021ME066) and the Qilu Young Scholar Program of Shandong University (to P.-F.X.), and the Taishan Scholars Project of Shandong Province (NO. tstp20230604).

## Notes

### Competing Interest Statement

The authors have declared no competing interest.

