## Supporting Information for "Programmable marine bacteria catalyze the valorization of lignin monomers"

* Correspondence:

Peng-Fei Xia

**Table S1.** Plasmids used in this study.

| Name | Description | Source |
|---|---|---|
| pQLL-mCherry | *pBR322* ori, *bla*, mCherry | Beijing Liuhe BGI. |
| pBBR1MCS-5 | *pBBR1* ori, *pBBR1* Rep, $Gm^R$ | Lab stock |
| pTemplate | *pUC* ori, *bla*, gRNA scaffold | Lab stock [1] |
| pCasEnv-lacI | pBBR1MCS-5, *lacI*-$P_{trc}$, *cas9* | Lab stock [2] |
| pWY06 | pWY, gRNA06 | Lab stock [2] |
| pgRNA-pcaG | pTemplate, gRNA07 | Lab stock [2] |
| pgRNA-pcaC | pTemplate, gRNA09 | This study |
| pmCherry | pBBR1MCS-5, *lacI*-$P_{trc}$, mCherry | This study |
| pWYi01 | pBBR1MCS-5, *lacI*-$P_{trc}$, *dcas9*, gRNA09 | This study |
| pWYi02 | pBBR1MCS-5, *lacI*-$P_{trc}$, *dcas9*, gRNA06 | This study |
| pWYi-M03 | pWYi02, gRNA09 | This study |
| pWYi-M04 | pWYi02, gRNA07 | This study |

**Table S2.** CRISPRi-mediated PCA production by *R. nubinhibens* at 36 h. [a]

| Target | Plasmid | IPTG (mM) | 4HB Consumption (mM) | PCA Titer (mM) | Yield (%, moL/moL) |
|---|---|---|---|---|---|
| / | pBBR1MCS-5 | 0.25 | 7.52 ± 0.39 | 0.33 ± 0.07 | 4.38 ± 1.11 |
| | | 0.5 | 9.69 ± 0.91 | 0.46 ± 0.43 | 5.00 ± 4.69 |
| *pcaC* | pWYi01 | 0.25 | 6.61 ± 0.43 | 0.39 ± 0.02 | 5.90 ± 0.12 |
| | | 0.5 | 8.28 ± 0.58 | 0.97 ± 0.03 | 11.74 ± 1.16 |
| *pcaH* | pWYi02 | 0.25 | 5.77 ± 0.76 | 0.54 ± 0.05 | 9.44 ± 0.30 |
| | | 0.5 | 7.26 ± 0.65 | 1.39 ± 0.06 | 19.27 ± 1.93 |

[a] Data are the averages and standard deviations from two independent biological experiments. IPTG, Isopropyl-β-D-thiogalactopyranoside; 4HB, 4-hydroxybenzoate; PCA, protocatechuate.

**Table S3.** Multiplex CRISPRi-mediated PCA production by *R. nubinhibens* at 36 h. [a]

| Target | Plasmid | 4HB Consumption (mM) | PCA Titer (mM) | Yield (%, moL/moL) |
|---|---|---|---|---|
| / | pBBR1MCS-5 | 9.69 ± 0.91 | 0.46 ± 0.43 | 5.00 ± 4.69 |
| *pcaC* and *pcaH* | pWYi-M03 | 7.41 ± 0.14 | 1 49 ± 0.12 | 20.15 ± 1.72 |
| *pcaH* and *pcaG* | pWYi-M04 | 5.88 ± 1.51 | 1.42 ± 0.22 | 24.47 ± 2.30 |

[a] Data are the averages and standard deviations from three independent biological experiments. IPTG, Isopropyl-β-D-thiogalactopyranoside; 4HB, 4-hydroxybenzoate; PCA, protocatechuate.

**Table S4.** Primers used in this study.

| Primer | Sequence |
|---|---|
| **Primers for In-Fusion DNA assembly** | |
| XIA-WY-109 | TTTCACACAGGAAACAGACCATGGTGAGCAAGGGCGAGGA |
| XIA-WY-110 | CGCTTACAATTTCCATTCGCCTACTTGTACAGCTCGTCCA |
| XIA-WY-111 | TCCTCGCCCTTGCTCACCATGGTCTGTTTCCTGTGTGAAA |
| XIA-WY-112 | TGGACGAGCTGTACAAGTAGGCGAATGGAAATTGTAAGCG |
| XIA-WY-093 | TTGACTACCGGAAGCAGTGTTCTAGATTGTAAAACGACGGCCAGTC |
| XIA-WY-094 | CATTTGAGAAGCACACGGTCACAGGAAACAGCTATGACCG |
| XIA-WY-095 | GACTGGCCGTCGTTTTACAATCTAGAACACTGCTTCCGGTAGTCAA |
| XIA-WY-096 | CGGTCATAGCTGTTTCCTGTGACCGTGTGCTTCTCAAATG |
| XIA-WY-201 | GATCATTTATTCTGCCTCCCTCGAACCACGCAATGCGTCT |
| XIA-WY-202 | CACCGTTTTTATCAGGCTCTTTGAGTGAGCTGATACCGCT |
| XIA-WY-203 | AGACGCATTGCGTGGTTCGAGGGAGGCAGAATAAATGATC |
| XIA-WY-204 | AGCGGTATCAGCTCACTCAAAGAGCCTGATAAAAACGGTG |
| **Primers for inverse PCR** | |
| XIA-WY-155 | TTTCAGACGCTGATCACCGAGTTTTAGAGCTAGAAATAGC |
| XIA-WY-156 | TCGGTGATCAGCGTCTGAAAGCTAGCATTATACCTAGGAC |
| XIA-WY-199 | TAGAGATCGCTAACTGGTTGGGACTGGTTGCATAACCATG |
| XIA-WY-200 | CAACCAGTTAGCGATCTCTAGTCACCTCCTAGCTGACTCA |
| **Primers for colony PCR** | |
| XIA-WY-044 | TTAGGTGGCGGTACTTGGGT |
| XIA-WY-045 | GCAGTCGCCCTAAAACAAAG |
| **Primers for RT-qPCR** | |
| XIA-WXY-001 | CCTGATCTAGCCATGCCG |
| XIA-WXY-002 | CGTATTACCGCGGCTGCT |
| XIA-WY-222 | TTTCGAGGAAATCCCCATGC |
| XIA-WY-223 | TTCCGCATAGGTTTCCTTGG |
| XIA-WY-224 | ATGACCAGGGCTATTACGTC |
| XIA-WY-225 | CGCCCTCGAAATAACATTGG |
| XIA-WY-226 | CATCGGTCTCAACACAAGGC |

| | |
|---|---|
| XIA-WY-227 | GGATGTCGAAACGGTAGACG |

**Table S5.** gRNA sequences used in this study.

| gRNA | Target | Strand [a] | PAM | Protospacer |
|---|---|---|---|---|
| gRNA06 | *pcaH* | C | CGG | CAAAGCGCCAGCGAAATCAC |
| gRNA07 | *pcaG* | C | AGG | GCAACGTCTCGACTACCTCA |
| gRNA09 | *pcaC* | C | AGG | TTTCAGACGCTGATCACCGA |

[a] C stands for coding strand and N stands for non-coding strand.

**Table S6.** Sequences of synthesized mCherry gene used in this study.

| **Gene** | **Sequence** [a] |
|---|---|
| mCherry | CGGTTCTGGCAAATATTCTGAAATGAGCTGTTGACAATTAATCATCC<br>GGCTCGTATAATGTGTGGAATTTCACACAGGAAACAGACCATGGTG<br>AGCAAGGGCGAGGAGGATAACATGGCCATCATCAAGGAGTTCATG<br>CGCTTCAAGGTGCACATGGAGGGCTCCGTGAACGGCCACGAGTTC<br>GAGATCGAGGGCGAGGGCGAGGGCCGCCCCTACGAGGGCACCCA<br>GACCGCCAAGCTGAAGGTGACCAAGGGTGGCCCCCTGCCCTTCGC<br>CTGGGACATCCTGTCCCCTCAGTTCATGTACGGCTCCAAGGCCTAC<br>GTGAAGCACCCCGCCGACATCCCCGACTACTTGAAGCTGTCCTTC<br>CCCGAGGGCTTCAAGTGGGAGCGCGTGATGAACTTCGAGGACGG<br>CGGCGTGGTGACCGTGACCCAGGACTCCTCCCTCCAGGACGGCG<br>AGTTCATCTACAAGGTGAAGCTGCGCGGCACCAACTTCCCCTCCGA<br>CGGCCCCGTAATGCAGAAGAAGACCATGGGCTGGGAGGCCTCCTC<br>CGAGCGGATGTACCCCGAGGACGGCGCCCTGAAGGGCGAGATCA<br>AGCAGAGGCTGAAGCTGAAGGACGGCGGCCACTACGACGCTGAG<br>GTCAAGACCACCTACAAGGCCAAGAAGCCCGTGCAGCTGCCCGGC<br>GCCTACAACGTCAACATCAAGTTGGACATCACCTCCCACAACGAGG<br>ACTACACCATCGTGGAACAGTACGAACGCGCCGAGGGCCGCCACT<br>CCACCGGCGGCATGGACGAGCTGTACAAGTAG [b] |

[a] Synthesized gene was carried by pQLL plasmid with $Amp^R$.

[b] The promoter sequence was shown in Blue and the coding sequence of mCherry was shown in Red.

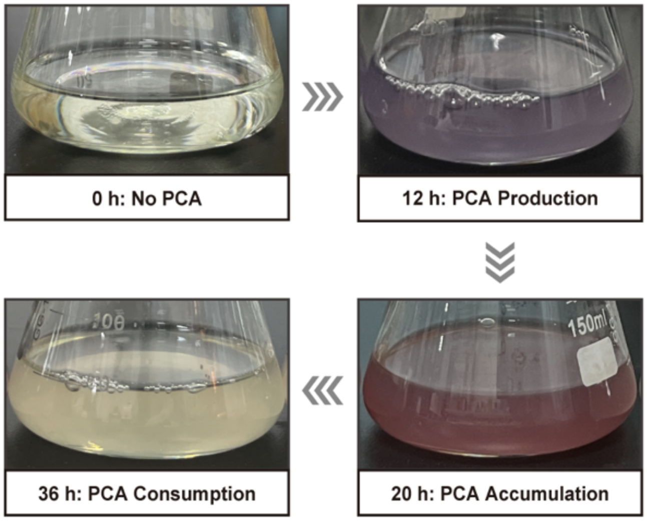


**Fig S1.** **The color variation of the wild-type *R. nubinhibens* during the cultivation in the basal medium with 4HB as the sole carbon source.**

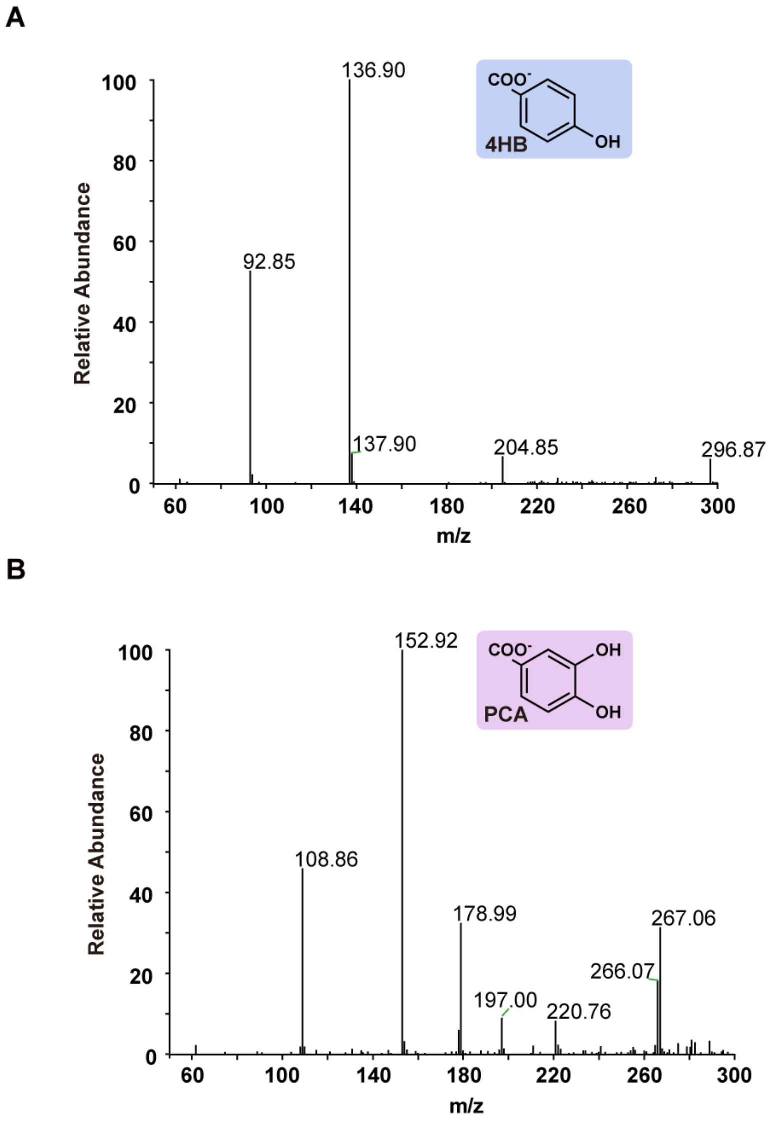


**Fig S2. MS analysis of standards. (A)** MS analysis of the 4HB standard. **(B)** MS analysis of the PCA standard.

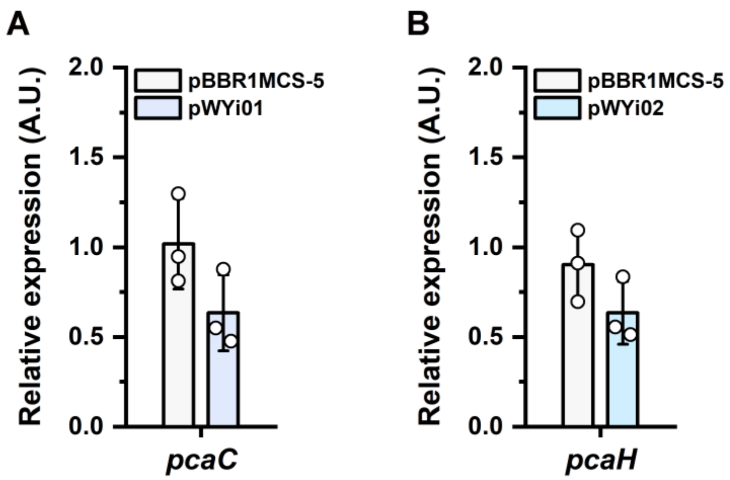


**Fig S3. Relative expression of target genes. (A)** Relative expression of *pcaC* in the strain with pWYi01. **(B)** Relative expression of *pcaH* in the strain with pWYi01. Experiments were carried in triplicate and the error bars represented the standard deviations of the means of three biological replicates.

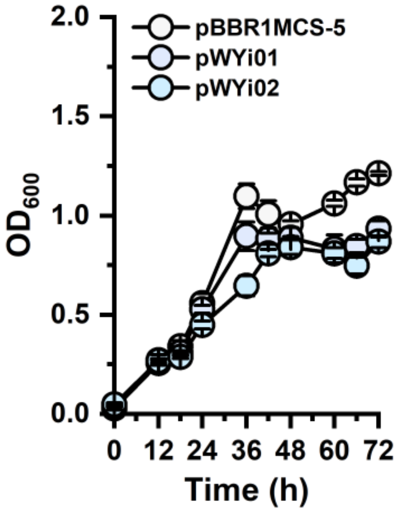


**Fig S4. Growth profile.** $OD_{600}$ of the strain with pBBR1MCS-5, pWYi01 and pWYi02 with 0.5 mM IPTG induction. Experiments were carried in triplicate and the error bars represented the standard deviations of the means of three biological replicates.

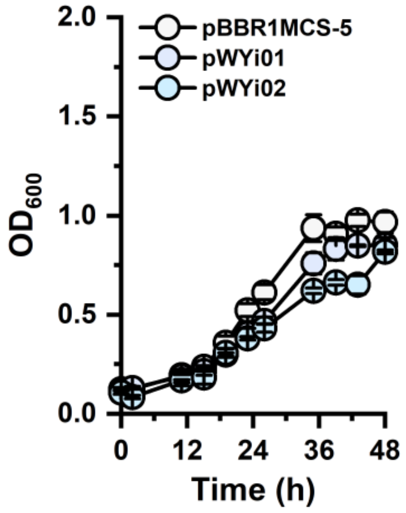


**Fig S5. Growth profile.** $OD_{600}$ of the strain with pBBR1MCS-5, pWYi01 and pWYi02 with 0.25 mM IPTG induction. Experiments were carried in triplicate and the error bars represented the standard deviations of the means of three biological replicates.

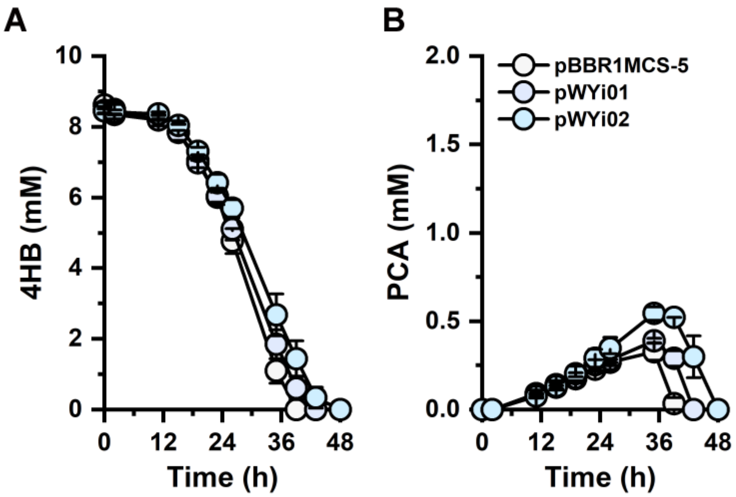


**Fig S6. The performance of PCA production in strains with pWYi. (A)** Utilization of 4HB, **(B)**Titer of PCA with pBBR1MCS-5, pWYi01 and pWYi02 with 0.25 mM IPTG induction. Experiments were carried out in duplicate and the error bars represented the standard deviations of the means of two biological replicates.

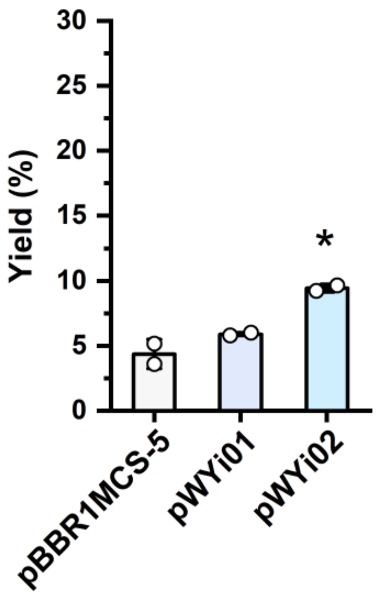


**Fig S7. Molar yield of PCA in strains with pWYi.** Yield (%) of PCA with pBBR1MCS-5, pWYi01 and pWYi02 with 0.25 mM IPTG induction. Experiments were carried out in duplicate and the error bars represented the standard deviations of the means of two biological replicates. The differences were statistically evaluated by t-test (*, $p<0.05$).

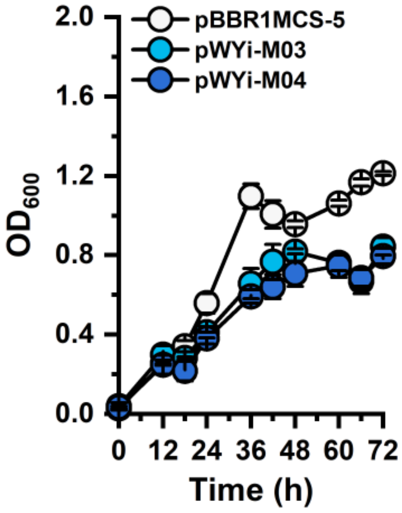


**Fig S8. Growth profile.** $OD_{600}$ of the strain with pBBR1MCS-5, pWYi-03 and pWYi-04 with 0.5 mM IPTG induction. Experiments were carried in triplicate and the error bars represented the standard deviations of the means of three biological replicates.
